# Interplay between mitochondria and diet mediates pathogen and stress resistance in *C. elegans*

**DOI:** 10.1101/154393

**Authors:** Alexey V. Revtovich, Ryan Lee, Natalia V. Kirienko

**Affiliations:** Department of BioSciences, Rice University, Houston TX, 77005, USA

**Keywords:** *C. elegans*, *E coli* HT115, OP50, BW25113, mitochondrial function, vitamin B12, mitochondrial homeostasis, pathogen sensitivity

## Abstract

Diet is a crucial determinant of organismal biology. Here we demonstrate the dramatic impact of a subtle shift in diet on the ability of *Caenorhabditis elegans* to survive pathogenic or abiotic stress. Interestingly, this shift occurs independently of canonical host defense pathways, arising instead from improvements in mitochondrial health. Using a variety of assays, we reveal that the most common *C. elegans* food source (*E. coli* OP50) results in a vitamin B12 deficiency that compromises mitochondrial homeostasis. Increasing B12 supply by feeding on *E. coli* HT115 or by supplementing bacterial media with methylcobalamin restored mitochondrial function, even if the bacteria were dead. B12 supplementation also efficiently increased host health without adversely affecting lifespan. Our study forges a molecular link between a dietary deficiency (nutrition/microbiota) and a physiological consequence (host sensitivity), using the host-microbiota-diet framework. The ubiquity of B12 deficiency (~10-40% of US adults) highlights the importance of our findings.

## Introduction

The diet and microbiota of an organism define its biology as much as its genome. Unfortunately, an avalanche of descriptive studies on model organism microbiomes have yielded very little mechanistic understanding of the relationships that define the host-microbiota-nutrition axis. One explanation for this is that the interplay amongst the three participants is incredibly complex and surprisingly dynamic. For example, the host and microbiota actively shape and, are in turn shaped by, their nutrition. Host genetics and environment determine initial susceptibility to microbial colonization, which will go on to influence all aspects of health. Establishing a mechanistic understanding of these interrelationships is crucial to the life history of an organism.

*Caenorhabditis elegans* offers a tantalizing system for simplifying these studies without sacrificing the ability to make discoveries that will be a useful starting point for more advanced organisms. At its most reduced form, the host-microbiota-nutrition axis of *C. elegans* can be collapsed to a binary system comprised of only two species, both of which are genetically tractable. As *C. elegans* is a bacteriovore, a single bacterial species can form the diet and the intestinal flora (due to incomplete disruption and digestion of bacterial food, which then colonizes the intestine). Generation of gnotobiotic worms is also simple and inexpensive.

Even this binary system has yielded significant insight regarding interactions between the host and its bacterial counterpart. For example, studies have shown that not only the relative biomacromolecular content of their diet (i.e., lipids, carbohydrates, etc.) but also bacterial metabolites informed the host’s biology, including folate (Virk et al., 2012), nitric oxide (Gusarov et al., 2013), and tryptophan (Anyanful et al., 2005; Gracida and Eckmann, 2013). Indeed, even the rate of bacterial respiration affected the metabolic state of the host (Saiki et al., 2008).

In this report, we show that differences in bacterial strains drive increased host sensitivity to a variety of stresses, including exposure to *P. aeruginosa*, oxidative stress, or hyperthermia. Through a panel of orthogonal assays, we demonstrated that a diet of *E. coli* strain OP50 causes a chronic vitamin B12 deficiency that perturbs mitochondrial health and function. B12 supplementation, even in the absence of living bacteria, increased resistance without shortening lifespan. Our findings provide a mechanistic explanation for the link between diet, cellular homeostasis, and organismal health.

## Results

### A diet of E. coli strain HT115 confers resistance to a variety of stresses in C. elegans

While characterizing a novel *C. elegans-P. aeruginosa* Liquid Killing assay (Conery et al., 2014; Kirienko et al., 2013), we made the unexpected observation that worms reared on *E. coli* HT115 exhibited increased resistance to *P. aeruginosa* compared to worms reared on *E. coli* OP50 (Fig 1A, B). This was true even if the bacteria lacked the L4440 RNAi vector. Pathogenesis in this assay requires the siderophore pyoverdine, which compromises host metabolism by removing ferric iron (Kirienko et al., 2015). Since exogenous iron strongly limits host killing (Kirienko et al., 2013), we tested whether worms reared on HT115 contained more iron, indirectly increasing their resistance to pyoverdine. However, neither a fluorometric iron (III) assay nor mass spectrometric measurement of total iron showed significant diet-dependent differences in host iron concentration (**Fig S1A, B**).

**Fig 1.**
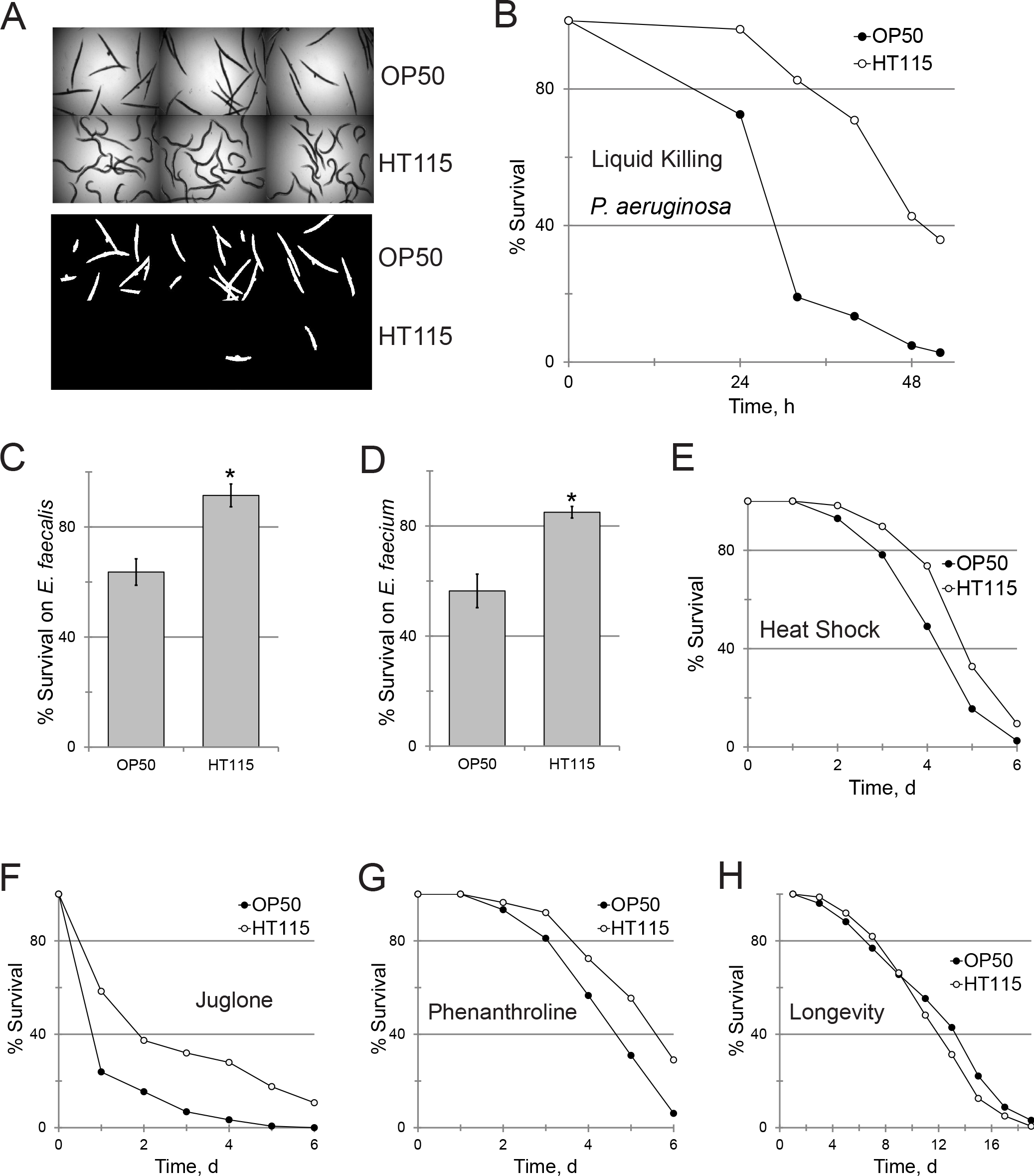
HT115 increases C. elegans’ resistance to stresses. **(A)** Representative brightfield (top) and fluorescent (bottom) images of *C. elegans* exposed to *P. aeruginosa* after feeding on OP50 or HT115. Sytox Orange, a cell-impermeant, fluorescent dye was used to mark dead worms. (**B**) Time course of quantification of staining for conditions in (**A**). (**C-G**) Survival of OP50- and HT115-fed worms after infection with *E. faecalis* or *E. faecium*, or exposure to thermal or oxidative stressors, or acute iron removal. (**H**) Lifespan of *C. elegans* fed with OP50 or HT115. *p*-value for (**B-G**) <0.01, (**H**)– n.s. See also Figures S1-3.

Substitution of *E. coli* OP50 with a variety of other bacterial foods (or UV- or heat-killed OP50) increases *C. elegans’* lifespan (Gomez et al., 2012; Kim, 2013; Win et al., 2013; Yu et al., 2015). These observations are typically interpreted to mean that OP50 is weakly pathogenic to *C. elegans* (Garigan et al., 2002; Garsin et al., 2001). On this basis, we surveyed basal gene expression levels for downstream effectors of a variety of *C. elegans* immune pathways, including PMK-1/p38, ZIP-2/bZIP, DAF-16/FOXO, FSHR-1/FSH, and SKN-1/Nrf2 (Estes et al., 2010; Garsin et al., 2003; Kim et al., 2004; Papp et al., 2012; Powell et al., 2009; Troemel et al., 2006). Worms reared on OP50 and HT115 had indistinguishable basal levels of expression of downstream innate immune effectors associated with these pathways (**Fig S2**).

Using either the conventional HT115 RNAi strain or an RNAi-competent OP50 derivative (Xiao et al., 2015), we knocked down key defense pathways and assayed sensitivity to *P. aeruginosa* exposure. Although some knockdowns affected the timing of death (e.g. *daf-2(RNAi)* prolonged survival, *daf-16(RNAi)* hastened death), worms fed HT115 survived longer in each case (**Fig S3**). These data indicate that weak OP50 pathogenicity (if it does exist) is unlikely to underlie the difference in sensitivity to *P. aeruginosa*.

Many defense pathways confer resistance to both abiotic and pathogenic factors. Therefore, we tested whether HT115 increased resistance to additional pathogens and abiotic stresses. As anticipated, survival was significantly increased when HT115-fed worms were infected with *Enterococcus faecalis* or *E. faecium* or were subjected to a heat shock or exposed juglone-induced oxidative stress or to the iron-scavenging xenobiotic 1,10-phenanthroline (Fig 1C-G). Increased resistance to pathogens and abiotic stresses is commonly associated with longer lifespan but, to our surprise, feeding HT115 did not extend lifespan (Fig 1H).

### Transcriptome profiling implicates mitochondrial defects in diet-induced stress sensitivity

To gain an unbiased snapshot of gene expression in worms fed OP50 or HT115, we performed transcriptome profiling. Interestingly, the number of differentially-regulated genes was very small, with only 35 genes upregulated between 2- and 8-fold in HT115 and 22 genes upregulated between 2- and 20-fold in OP50 (**Table S1**). Strikingly, of the latter 22 genes, 12 encoded proteins localized to the mitochondria (Table 1). This enrichment was highly significant (*p*=2.2*10^-16^), particularly given that the fraction of nuclear genes that encode mitochondrial proteins in *C. elegans* and humans is ~6% and 7% respectively (Li et al., 2009; Prokisch et al., 2006). This enrichment was specific to genes upregulated by an OP50 diet; amongst genes expressed at higher levels in HT115-fed worms, only two mitochondrial genes (*acox-2* and T22B7.7) were observed (2/35=5.7%, *p*=0.9896). These data argue that feeding with OP50 increases expression of mitochondrial genes. Two genes in particular caught our attention: *hsp-60*, which encodes a mitochondrial chaperone expressed constitutively at low levels but induced upon mitochondrial damage (Benedetti et al., 2006; Haynes et al., 2007; Yoneda et al., 2004), and *acdh-1*, which encodes a short-chain acyl-CoA dehydrogenase that has been used as an indicator of branched chain amino acids and/or propionyl-CoA, a mitotoxic byproduct of their metabolism (Schwab et al., 2006; Watson et al., 2014).

**Table 1.**
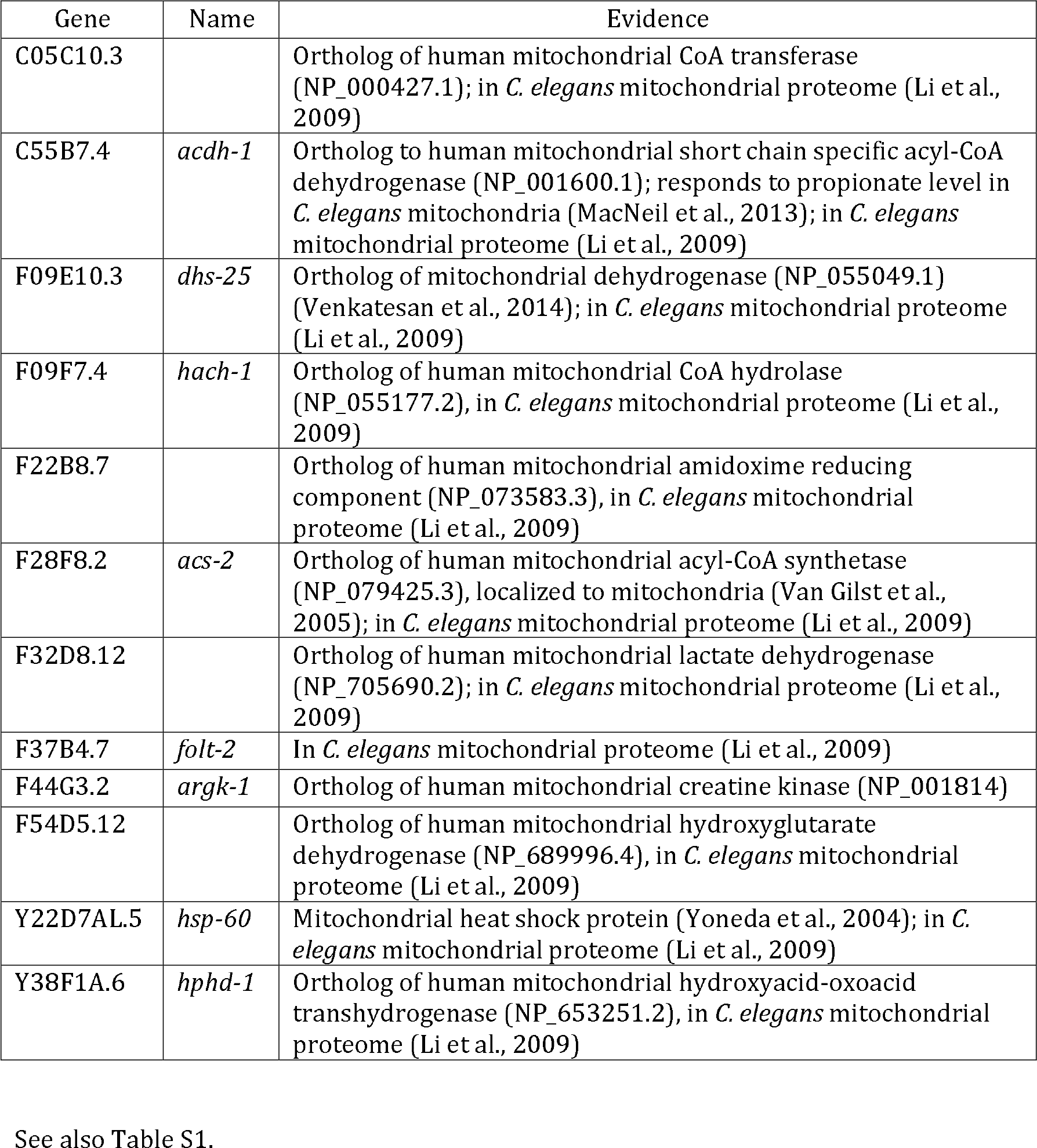
Mitochondrial genes upregulated by feeding on E. coli OP50.

### A diet of HT115 improves the vitamin B12 deficiency caused by OP50

Consistent with our microarray results, we observed increased *acdh-1::*GFP expression in OP50-fed worms, whether measured by conventional imaging (Fig 2A,B) or by COPAS flow vermimetry (Fig 2C). Vitamin B12 is an essential micronutrient for most organisms, including *C. elegans* and *E. coli* and is required for efficient metabolism of branched chain amino acids and propionyl-CoA (Watson et al., 2014; Watson et al., 2015). Our data, and those of the Walhout lab, demonstrate that an OP50 diet induces a B12 deficiency that triggers compensatory increases in expression of *acdh-1::*GFP and other mitochondria-related genes (Watson et al., 2016).

**Fig 2.**
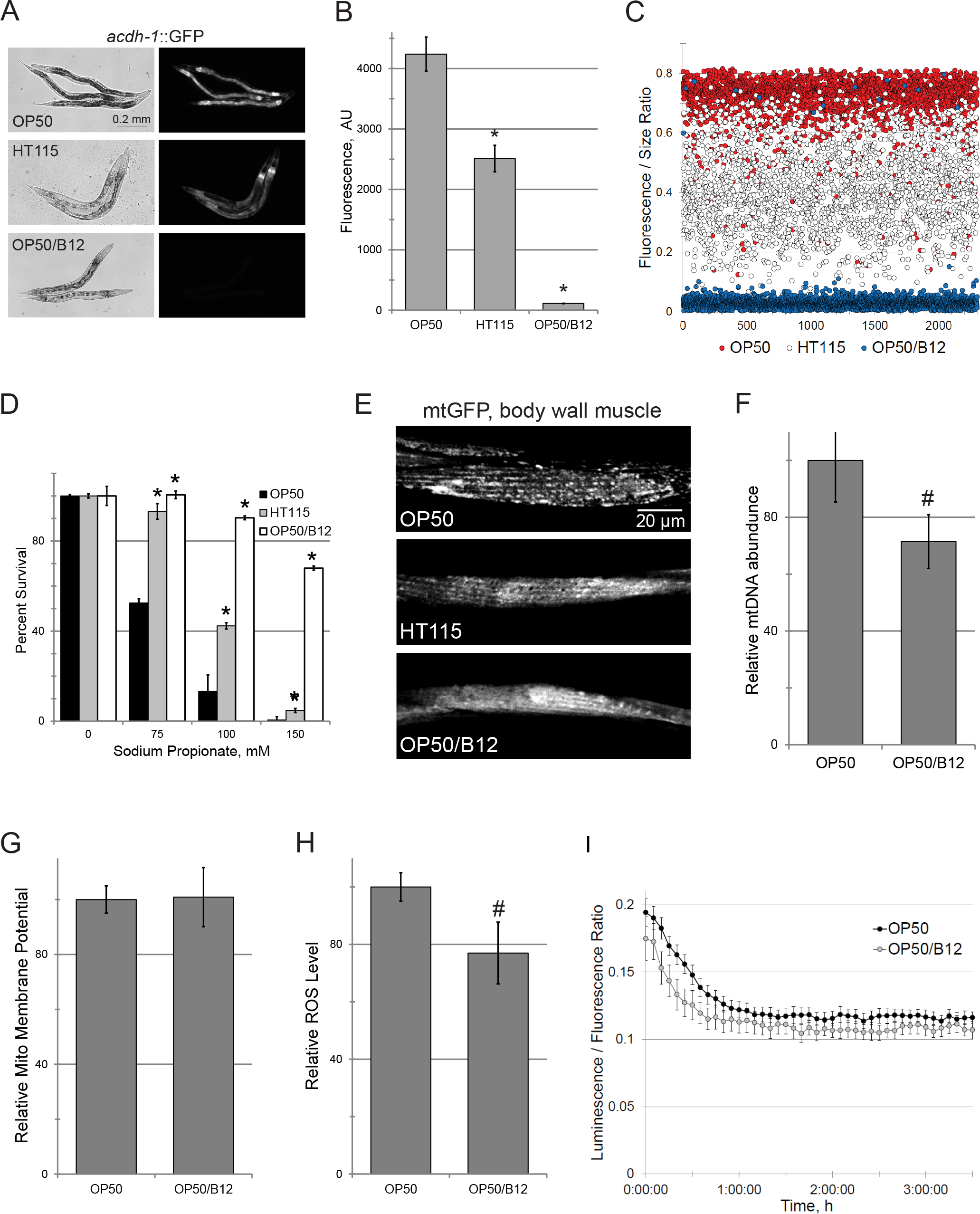
HT115 or methylcobalamin supplementation improve mitochondrial health. (**A-C**) Visualization and quantification of *acdh-1::*GFP reporter fluorescence in worms fed various diets, as indicated. (**D-E**) Effect of the diet on propionate toxicity (**D**) and connectivity of mitochondrial network (**E**). (**F-I**) Impact of methylcobalamin (B12) supplementation on mitochondrial count (F), membrane potential (**G**), and production of ROS (**H**) or ATP (**I**). * - *p*<0.01, # - *p*<0.05

We attempted to use mass spectrometry to measure S-adenosylcobalamin in worms fed either HT115 or OP50 (or in the bacteria themselves) to obtain quantitative measurements of vitamin B12. Unfortunately, B12 levels for all samples measured were below the detection threshold, even when samples were concentrated (*data not shown*). To circumvent this limitation we used sodium propionate sensitivity as a functional readout of B12 sufficiency. Consistent with other data, a diet of HT115 significantly increased resistance to propionate, indicating that it improves the B12 shortage (Fig 2D).

### Vitamin B12 deficiency disrupts mitochondrial homeostasis

We hypothesized that the dietary B12 deficiency causes buildup of propionyl-CoA, which would lead to mitochondrial damage. Therefore, we assayed mitochondrial homeostasis in *C. elegans* fed either OP50 or HT115. Under normal conditions, mitochondrial quality control involves constant fission and fusion events that serve to pool healthy, functional mitochondrial content, while damaged material is sequestered for autophagic recycling (Liesa and Shirihai, 2013). Compared to mammals, *C. elegans* mitochondria generally exhibit greater interconnectivity and longer tubular architecture (e.g., disrupting *C. elegans* mitochondrial genes almost always increases fragmentation (Ichishita et al., 2008)). We used a *C. elegans* strain expressing a mitochondrially-targeted GFP (Benedetti et al., 2006) to assay mitochondrial health after OP50 or HT115 consumption. Worms fed HT115 demonstrated increased connectivity and less clumping (i.e., punctae) than OP50 (Fig 2E), indicating improved mitochondrial health.

### Vitamin B12 supplementation improves mitochondrial health

To determine whether we could remedy the B12 limitation, we spiked OP50 growth medium with exogenous methylcobalamin to a final concentration of 0.2 µg/mL. Supplementation dramatically decreased *acdh-1::*GFP fluorescence (Fig 2A-C), increased resistance to propionate (Fig 2D), and improved mitochondrial network architecture (Fig 2E). Combined, these data suggested that methylcobalamin supplementation significantly improved mitochondrial health.

We assayed other metrics of mitochondrial health, including mitochondrial count (Fig 2F), membrane potential (Fig 2G), production of reactive oxygen species (ROS, Fig 2H), and ATP (Fig 2I) to verify this conclusion. Interestingly, methylcobalamin supplementation decreased the number of mitochondria while mitochondrial membrane potential remained steady, indicating that the average membrane potential of each mitochondrion was slightly increased by treatment. Methylcobalamin also significantly decreased ROS production, while ATP production decreased only slightly. The most parsimonious explanation for these observations is that a dietary B12 deficiency significantly compromises mitochondrial health and efficiency.

### Vitamin B12 deficiency in OP50 diet drives sensitivity to stress

We predicted that the mild mitochondrial dysfunction exhibited by OP50-fed worms may drive their increased sensitivity to stress. We reared worms on OP50 or OP50 supplemented with methylcobalamin (OP50/B12), and tested their resistance to *P. aeruginosa*. We observed a dramatic difference in host survival: virtually all OP50-fed worms were dead by the time OP50/B12-fed worms showed ~10% death (Fig 3A, B). Bacterial metabolism of B12 was superfluous for this effect; *C. elegans* fed heat-killed *E. coli* spotted onto plates containing methylcobalamin-supplemented media also exhibited virtually no *acdh-1::*GFP expression and increased survival when exposed to *P. aeruginosa* (**Figs S4A, 3C**). Cobalamin supplementation also substantially improved survival after infection with *E. faecalis* or *E. faecium* (Fig 3D, E). Next, we examined whether cobalamin was beneficial during exposure to more generic stressors, like hyperthermia. Supplementation again significantly increased *C. elegans* survival (Fig 3F).

**Fig 3.**
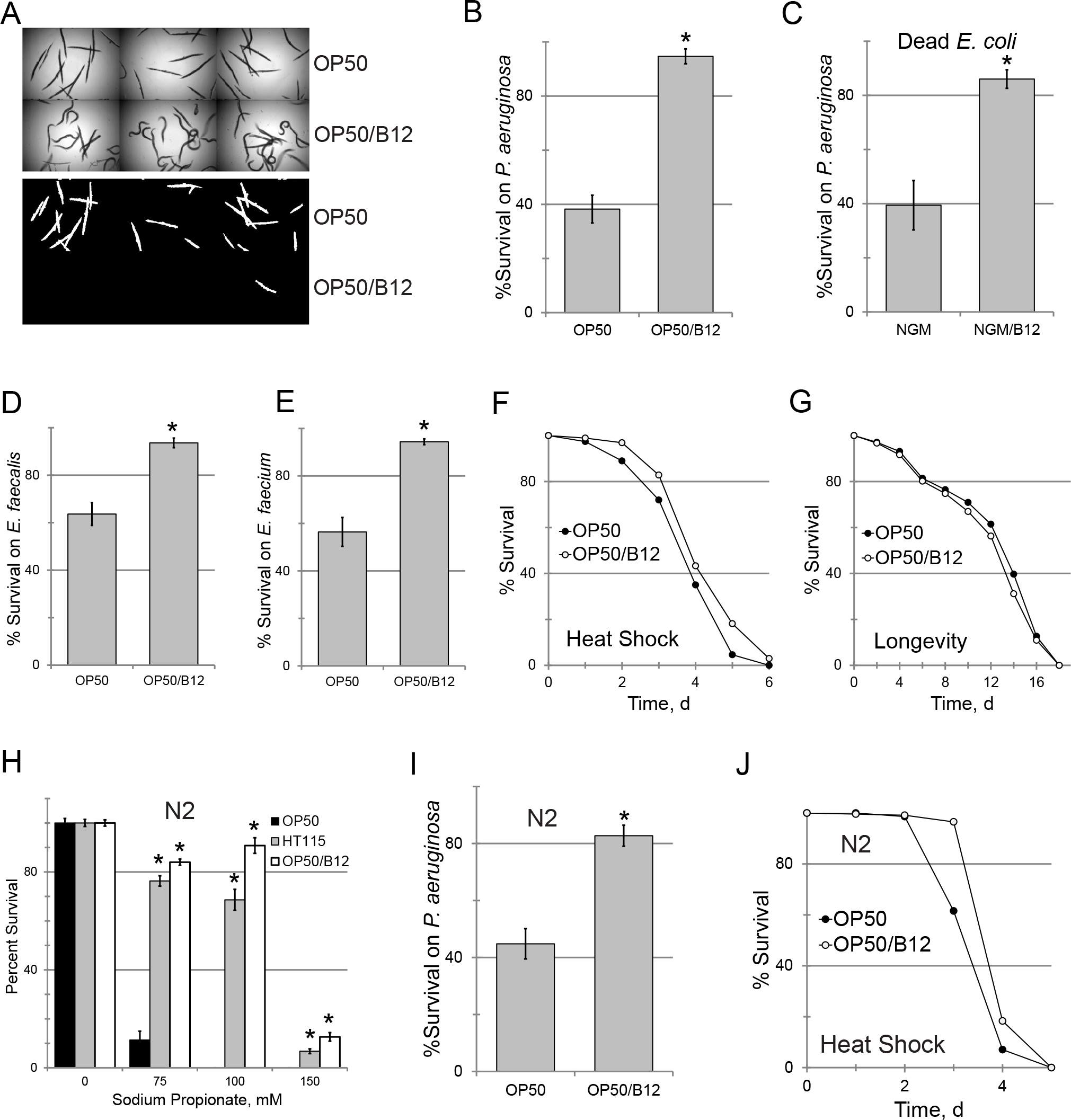
Methylcobalamin supplementation increases resistance to stress. (**A-B**) Representative images and quantification of survival of *C. elegans* exposed to *P. aeruginosa* after feeding on OP50 with or without methylcobalamin (B12) supplementation. Fraction of surviving worms here and elsewhere was inferred based on staining with Sytox Orange, a cell impermeant dye. (**C**) Relative survival of *C. elegans*, grown on NGM or NGM/methylcobalamin plates with heat-killed OP50 as the food source, after exposure to *P. aeruginosa*. (**D-G**) Effect of methylcobalamin-supplemented OP50 feeding on *C. elegans* resistance to infection with *E. faecalis*, *E. faecium*, heat shock, or lifespan. (**H-J**) Survival of wild-type worms fed OP50 with or without methylcobalamin supplementation after exposure to propionate (**H**), *P. aeruginosa* (**I**), or hyperthermia (**J**). * - *p*<0.01 (**B-E, H, I**). See also Figure S4.

A growing body of evidence suggests that decreasing mitochondrial activity, either by genetically compromising oxidative phosphorylation or by caloric restriction, extends lifespan (Luz et al., 2016; Yang and Hekimi, 2010). Since B12 supplementation improves mitochondrial function, we wanted to know whether it may be beneficial during adverse conditions, but would also ultimately shorten lifespan. No significant changes in lifespan were seen in worms receiving a diet supplemented with methylcobalamin (Fig 3G).

All of the experiments described above were performed in a *glp-4(bn2ts)* background, which fails to generate a germline and is sterile at non-permissive temperatures. Sterility is necessary to avoid non-specific death in the Liquid-Killing assay. To rule out the possibility that the observed phenotypes were a result of the *glp-4 (bn2)* background, we tested wild-type N2 worms. In every case tested, wild-type worms recapitulated our findings from *glp-4(bn2)* mutants (Fig 3H-J).

### The E. coli TonB transporter mediates vitamin B12 internalization and subsequent host health

Since *E. coli* is unable to synthesize vitamin B12, it must be imported from the extracellular milieu. Efficient import has been reported to require TonB activity (Bassford et al., 1976; Kadner, 1990), so we evaluated *acdh-1::*GFP fluorescence in worms fed *E. coli* with deletions of the *tonB* gene (Baba et al., 2006). While *acdh-1::*GFP fluorescence in worms reared on the parental strain (*E. coli* BW25113) was similar to OP50 (**Fig S4B**), tonB deletion slightly, but reproducibly, increased GFP level (**Fig S4C**, p<0.01), consistent with a previous report (Watson et al., 2014). Correspondingly, *tonB* deletion also increased sensitivity to *P. aeruginosa* exposure (**Fig S4D**). As with OP50, supplementation of wild-type BW25113 with methylcobalamin completely abolished *acdh-1::*GFP fluorescence (**Fig S4E**). In contrast, tonB mutants showed only partial attenuation of GFP expression after methylcobalamin supplementation, indicating that the bacteria were unable to efficiently internalize the vitamin (**Fig S4F, G**). One possible explanation for the attenuation in GFP fluorescence that we observed in these mutants is that *E. coli* has a salvage pathway to acquire precursors for B12 biosynthesis (Fang et al., 2017). Our observation that *tonB* mutation increases host sensitivity to stress still further (**Fig S4D**) indicates a role for TonB activity in the health of the host.

## Discussion

Recently, Leulier and colleagues defined the “nutrient-microbiota-host axis” to incorporate all of these determinants into a single conceptual framework (Leulier et al., 2017). Our results leverage this idea to connect diverse phenomena into an articulated whole. For example, our data indicate that a dietary deficiency in vitamin B12 deficiency causes mitochondrial defects, likely through the buildup of propionyl-CoA (a known mitochondrial toxin). We also demonstrated that this damage significantly increased sensitivity to a variety of stresses. Recently, Lin and Wang demonstrated that methionine supplementation alters the metabolism of the food-microbiota of *C. elegans*, directly influencing host mitochondrial health (Lin and Wang, 2017). Interestingly, methionine supplementation did not influence host survival after exposure to P. aeruginosa in Liquid Killing conditions (56.5% vs 54.2%, *p*=0.363).

Our findings provide a mechanistic explanation for clinical observations that patients with methylmalonic or propionic acidemias (either of which can arise from vitamin B12 deficiency) exhibit symptoms that are strikingly similar to those with congenital mitochondrial defects (including poor growth, muscle weakness, and disorders of the liver, kidney, and gastrointestinal and respiratory tracts). It also explains why mice with inactive methylmalonyl-CoA mutase (one of two B12-dependent enzymes) exhibit reduced respiratory chain activity (Chandler et al., 2009). Vitamin B12 is clearly important for normal cellular stress resistance and healthy aging. Despite this, estimates of deficiency range between 10-40% of the population, with increased prevalence in the elderly (Spence, 2016).

Previous studies from the Walhout lab showed that some diets replete in B12 (including *Comamonas aquaticus* and *E. coli* HT115) were associated with shorter lifespans, although the difference is not directly due to B12 (MacNeil et al., 2013). We also saw a small, but statistically significant, decrease in maximum lifespan when *C. elegans* was reared on *E. coli* HT115. Intriguingly, methylcobalamin did not affect the lifespan of worms when it was added to an OP50 diet (Fig 3E), suggesting that the improvement in mitochondrial health provided by B12 supplementation can be unlinked from lifespan shortening. Currently, we hypothesize that some other aspect of *Comamonas* and HT115 diets, probably a common metabolite, was responsible for this effect.

Interestingly, even though we started to query the host-microbiota-diet axis with a minimal, binary system (*C. elegans* and *E. coli*), a number of unexpected complexities have arisen. For example, subtle differences in the biology of *C. elegans* reared on OP50 or HT115 have generally been attributed to the differences in their origins (i.e., OP50 was derived from an E. coli B strain while HT115 came from *E. coli* K12). However, BW25113 (which also originated from K12) showed *acdh-1::*GFP fluorescence similar to that of OP50. Heat-killed OP50 showed *acdh-1::*GFP fluorescence similar to that of HT115-fed *C. elegans*. These data indicate that *acdh-1::*GFP expression and mitochondrial health poorly correlate with strain origin and are likely to be dynamic, illustrating the subtle nuances that lie along the nutrient/microbiota axis and the dramatic impacts they have on host phenotype.

We also showed that this framework can help identify an important mechanism used by *C. elegans*, and presumably more complex metazoans, to fine-tune their cell metabolism to resist adverse environmental factors (including pathogens, heat, and toxins), even without activating canonical host defense pathways. Interestingly, the sensitivity to stress caused by B12 deficiency seems to correlate with the degree of mitochondrial involvement in the stress-induced pathology. For example, we observed that methylcobalamin supplementation decreased *acdh-1* expression by 10-fold or more while proportional increases were seen in resistance to propionate or *P. aeruginosa* in Liquid Killing. This result corroborates our recent findings that place mitochondrial homeostasis at the heart of this pathogenesis model (Tjahjono and Kirienko, 2017). Although the effect was smaller, benefits from B12 supplementation were still observed when stresses were more pleiotropic (such as heat shock or oxidative stress, where intracellular contents are more uniformly damaged). Taken together, our findings emphasize the importance of using a bottom-up approach to effectively understand the mechanisms that connect nutrition, metabolic activity, and the host’s microbiota to its health.

## Experimental Procedures

Additional detailed methods are presented in the Supplemental Experimental Procedures available online.

### C. elegans pathogenesis and stress assays

Liquid- and Slow-Killing assays were performed as described elsewhere (Kirienko et al., 2014). Killing assays using *Enterococcus faecalis* strain OG1RF and *E. faecium* strain E007 assays were performed similarly, except that *P. aeruginosa* strain PA14 was substituted with the relevant pathogen, 10% brain-heart infusion medium was substituted for 20% slow-kill media, and host death was measured after 4 days of infection. Propionate toxicity was measured as described (Watson et al., 2014). For oxidative and iron removal stresses, NGM agar plates were supplemented with 120 µM of juglone (Kirienko and Fay, 2010) or 100 µM of phenanthroline, respectively. Heat shock experiments were carried out at 30 °C, lifespan was measured at 25 °C. Wild-type *C. elegans* were rendered sterile by *cdc-25.1(RNAi)* when appropriate. For agar-based assays, three plates with 50 worms/plate/strain/biological replicate were used. For liquid-based experiments, 10 wells with 20 worms/well/strain/biological replicate were used. At least three biological replicates were performed for each experiment. Statistical significance was calculated based on log-rank test, except Student’s t-test was used for Liquid Killing and propionate toxicity.

## Author Contributions

AVR, RL, and NVK conceived of and performed experiments. AVR and NK analyzed data and wrote the manuscript. NK edited the manuscript and supervised the work.

## Acknowledgements

We wish to thank Alex Kang and Dr. Chris Pennington for technical assistance with ICP-MS. *E. coli* OP50(xu363) was provided by Dr Eyleen O’Rourke, *E. faecalis* and *E. faecium* strains were the kind gift of Dr. Danielle Garsin; some strains were provided by the CGC, which is funded by NIH Office of Research Infrastructure Programs (P40 OD010440). This study was supported by the Cancer Prevention and Research Institute of Texas (CPRIT) RR150044 and National Institutes of Health K22 AI110552 awarded to NVK.

